# Unsupervised pattern discovery in spatial gene expression atlas reveals mouse brain regions beyond established ontology

**DOI:** 10.1101/2023.03.10.531984

**Authors:** Robert Cahill, Yu Wang, R. Patrick Xian, Alex J. Lee, Hongkui Zeng, Bin Yu, Bosiljka Tasic, Reza Abbasi-Asl

## Abstract

The rapid growth of large-scale spatial gene expression data demands efficient and reliable computational tools to extract major trends of gene expression in their native spatial context. Here, we used stability-driven unsupervised learning (i.e. staNMF) to identify principal patterns (PPs) of 3D gene expression profiles and understand spatial gene distribution and anatomical localization at the whole mouse brain level. Our subsequent spatial correlation analysis systematically compared the PPs to known anatomical regions and ontology from the Allen Mouse Brain Atlas using spatial neighborhoods. We demonstrate that our stable and spatially coherent PPs, whose linear combinations accurately approximate the spatial gene data, are highly correlated with combinations of expert-annotated brain regions. These PPs yield a new brain ontology based purely on spatial gene expression. Our PP identification approach outperforms principal component analysis (PCA) and typical clustering algorithms on the same task. Moreover, we show that the stable PPs reveal marked regional imbalance of brainwide genetic architecture, leading to region-specific marker genes and gene co-expression networks. Our findings highlight the advantages of stability-driven machine learning for plausible biological discovery from dense spatial gene expression data that are infeasible by conventional manual approaches.

## Introduction

In the past decade, unsupervised explorations of large-scale single-cell transcriptomics datasets enabled by machine learning tools led to an unbiased definition of cell types––groups of cells with similar gene expression patterns^1–5^. Traditionally, genetic profiling requires cell isolation that discards the spatial information of cells within tissues or organs. Spatially resolved techniques preserve the spatial information which are crucial for understanding cell function and tissue organization^6–8^. Spatial patterns may correlate with specific cell types or cell type combinations and reflect local tissue characteristics in structure and function. To accommodate their growing popularity and data throughput, computational pipelines also need to incorporate spatial information in interpreting the outcome. Spatially-aware analytical tools apply to both healthy and diseased tissues and may help elucidate gene and organ function and generate viable hypotheses for disease mechanisms^9–13^ in the spatial domain.

A core element of biospatial information is the anatomical atlas of an organ, which is defined by expert annotation based on accumulated historical data. Brain atlases^14,15^ are comparable across individuals and species, facilitating cross-referencing and analysis of neural data in a consistent manner. For the adult mouse brain, the Allen Common Coordinate Framework (CCF 3.0) is a widely used atlas and ontology (hierarchical relations between parts of the atlas) built on the Allen Mouse Brain Atlas (ABA)^16^. Yet its construction is time-intensive, hard-to-scale, and potentially affected by human judgment. Data-driven approaches can mitigate human error, streamline the process, and uncover information hard to perceive by the human eye^17^.

Currently, segmentation and clustering are the two main categories of machine learning approaches in the analysis of spatial gene expression data^17–31^. While these methods yield a set of spatially non-overlapping or in some cases overlapping regions, the problem formulation focuses on local information and does not explicitly model the global structure of the entire gene expression data. By contrast, matrix decomposition techniques such as non-negative matrix factorization (NMF)^32–34^ provide a model-based representation of an entire dataset as a combination of a set of dictionary elements or principal patterns (PPs)^18,35–40^. These models could reduce complex spatial patterns into a combination of PPs, which provide a more interpretable representation of each data point compared to segmentation or clustering. However, the simplest matrix decomposition model, principal component analysis (PCA), despite its frequent usage^41–43^, is not a sensible choice because biologically realistic assumptions, such as non-negativity, are unmet. NMF and its variants include non-negativity as an explicit constraint in the problem formulation, leading to a more biologically plausible outcome^36^, with relevant applications in the analysis of gene expression^18,35,37,44^, neural recordings^38,39^, etc.

More importantly, we used stability-driven NMF (staNMF) algorithm^18^ to incorporate stability as the central criterion in model selection to analyze spatial gene expression datasets of the adult mouse brain. Stability is a measure of scientific reproducibility and statistical robustness^45^. It asks whether each step of the pipeline produces consistent results with slight perturbations in the model or data^46,47^. It is a minimum requirement for interpretability^45,47^ and, in the current context, essential for identifying biologically meaningful and coherent spatial patterns in the mouse brain^48^. Previous work has demonstrated the promise of staNMF in interpreting 2D spatial gene expression images from *Drosophila* embryos^18^. Here, we extend the analysis to 3D and, by spatial correlation analysis with an existing brain atlas^16^, discover that the PPs are clearly localized in single or combinations of anatomical regions, which suggests a gene expression-defined ontology beyond the one from neuroanatomy. Moreover, our analysis reveals the marked regional imbalance of gene expression, with the hippocampus having the most diverse gene expression than others, followed by the isocortex and the cerebellar regions. We recover the spatial genetic architecture using the spatial organization and correlation structure of the gene expression, which reveal region-specific marker genes as well as putative spatial gene co-expression networks (sGCNs) spanning the entire mouse brain.

## Results

### Identifying stable PPs in the Allen Mouse Brain Atlas

Our computational pipeline is designed to extract PPs in the spatial gene expression data with additional pre- and post-processing steps for data preparation, quality assessment, and to derive biological insights (**Fig. 1A**). PPs or the latent factors that optimally capture data variability are extracted using staNMF^13,30^, which involves stability analysis to ensure the reproducibility of the PPs (referred to as stable PPs). The analysis evaluates an instability score, here defined as the average dissimilarity of all learned dictionary pairs using their cross-correlation matrix. We use the Hungarian matching method^49^ (**Fig. 1B**) or an Amari-type error function^50^ (**Fig. S1**) to account for the invariances (see **Methods)**. Overall, staNMF yields two outputs: (1) *K* PPs for the whole imaging data and (2) the coefficients or PP weights for each gene expression image. The model reconstructs each 3D gene expression profile by a non-negative linear combination of the PPs. Each PP is calculated after model training as one of *K* dictionary elements learned via staNMF. The weights of a PP or dictionary element for each gene are determined by the coefficients of staNMF. Our end-to-end pipeline is computationally efficient and can handle large datasets generated in modern spatially resolved sequencing techniques^8,48^.

**Figure 1:**
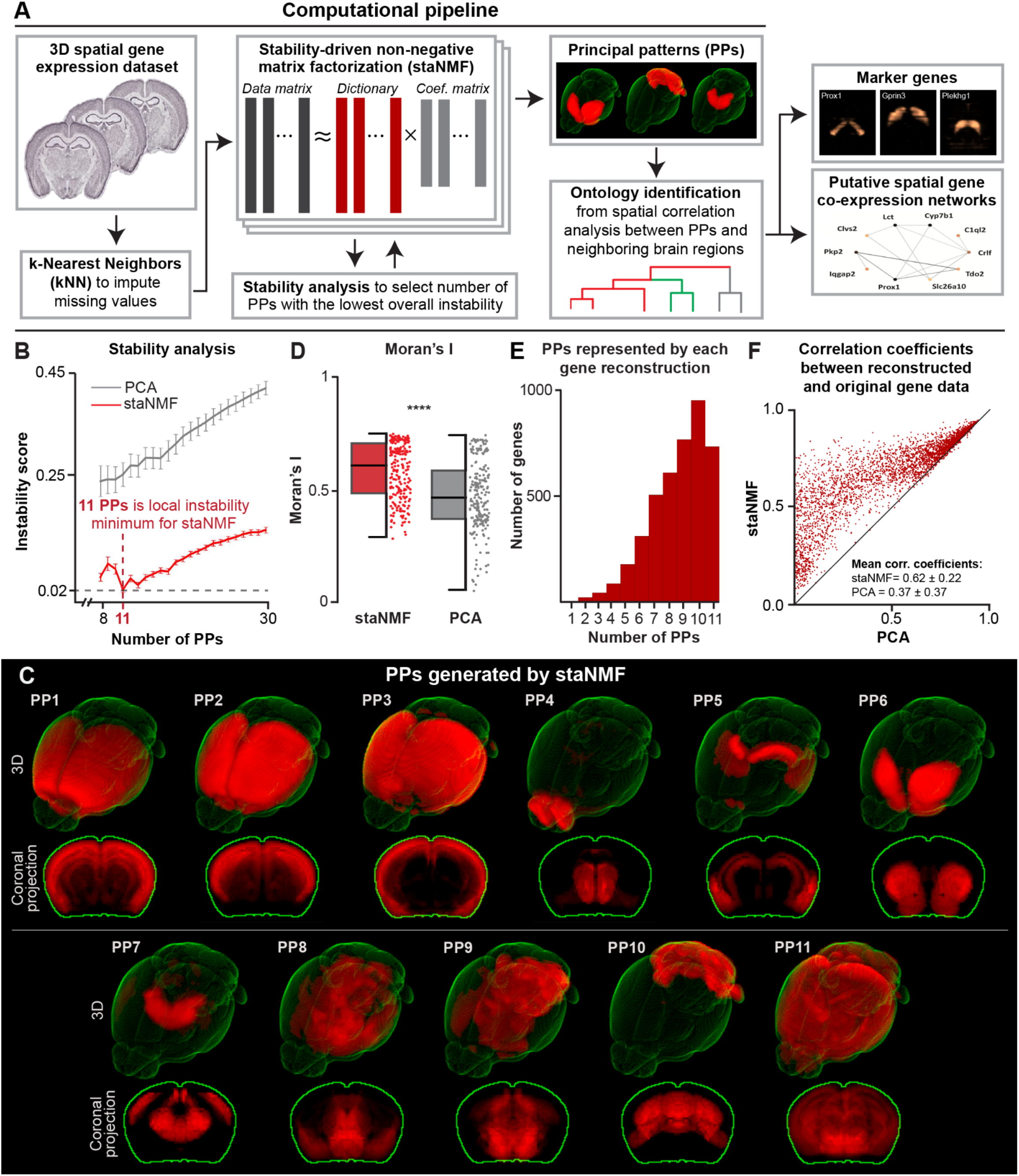
staNMF-based computational pipeline for spatial gene expression data. **A**. Illustration of the computational pipeline with essential steps and outcomes. **B**. Stability analysis for staNMF PPs and PCA PPs across 100 runs for each *K* value, from 8 to 30 for ABA dataset, using the Hungarian matching method. Error bars are the standard deviation. **C**. 11 PPs generated by staNMF from the ABA dataset in 3D and projected on the coronal plane. **D**. Boxplot of Moran’s I calculated for staNMF vs. PCA PPs across 220 bootstrapped simulations (p-value<0.001). The data from each individual point is shown in a vertical column to the right of the boxplot. **E**. The number of PPs represented by each staNMF gene reconstruction of the 4,345 ABA genes. **F**. Comparison of the reconstruction accuracy between staNMF and PCA. Each dot represents one gene. The coordinates of each dot are the Pearson correlation coefficients between the measured gene expression in the ABA dataset and the reconstructed gene expression by staNMF or PCA.

We used the pipeline to determine the PPs for 4,345 3D spatial gene expression profiles in the adult mouse brain (56 days old) from the ABA dataset^48^, where each gene was examined by whole-brain serial sectioning and RNA *in situ* hybridization (ISH) at 200 μm isotropic resolution. The preprocessing step uses a kNN-based voxel imputation to fill in approximately 10% missing voxel data in ABA. On a hold-out test set of 1,000 random voxels for each of the 4,345 genes from the ABA dataset (for 4,435,000 total hold-out data points), the mean error was smaller than 0.01. The Pearson correlation coefficient (PCC) between the measured and imputed gene expression data was 0.52, with a p-value < 0.01. To determine stable PPs, we calculated the instability score with 100 runs of the same algorithm across a range of 8 to 30 possible numbers of PPs. The lowest instability (and thus highest stability) was found when *K*=11, with an instability score of 0.020 ± 0.002 (1 is the maximum instability) for the Hungarian matching method^49^. The settings when *K*=13 and *K*=12 have the next two lowest instability scores (0.03 and 0.04, respectively, with standard deviations < 0.01).

To assess the performance of our approach, we first compared the outcome of staNMF with PCA (**Fig. 1B**). Our results show that staNMF PPs have higher stability and lower standard deviation vs. PCA PPs (0.25 ± 0.01) at every value of *K* tested. In terms of computational runtime, staNMF takes longer to run than PCA, though both are fast-running models. On a 2021 MacBook Pro M1 laptop CPU, where the computation was tested, it takes 26 seconds to run staNMF to create one set of PPs on the ABA dataset vs. 4 seconds for PCA.

Another important aspect to evaluate is the spatial coherence of PPs, which is important for their biological interpretability. To this end, we used Moran’s I^42,51–53^, which ranges in value from –1 to 1, as a global summary statistic (**Fig. 1C**). A value close to -1 indicates little spatial organization, whereas a value close to 1 indicates a clear spatially distinct pattern. The average Moran’s I for staNMF PPs is 0.58 ± 0.12, which is considerably higher than that of PCA at 0.47 ± 0.15 (p-value<0.001) across 20 bootstrapped simulations for each of the 11 PPs (**Fig. 1D** and **Fig. S2**). This suggests a stronger spatial separation and coherence of PPs obtained from staNMF than those from PCA (See **Fig. S3A** for visualization of PCA PPs). We want to point out that although staNMF PPs are spatially coherent, a large number of PPs tend to be present in most gene expression profiles (58% of all genes are represented in 9 or more PPs), suggesting the spatial heterogeneity of gene expression in the adult mouse brain (**Fig. 1E**). Only two genes are represented in a single PP (<0.1% of all 4,345 genes), while 438 genes are represented in all 11 PPs (10.1% of all genes).

Additionally, we compared the staNMF and PCA reconstruction accuracy in a scatterplot (**Fig. 1F**), where each point represents one of the 4,345 genes in the dataset. We defined the reconstruction accuracy as the PCC between the reconstructed and the original gene expression image. **Fig. 1F** shows that staNMF considerably outperforms PCA in the reconstruction performance (0.62 ± 0.22 for staNMF compared to 0.37 ± 0.37 for PCA; 24% higher accuracy for staNMF). We also found that our kNN imputation of missing values improves staNMF’s reconstruction accuracy of the original data set from 0.59 to 0.62. It is worth noting that the reconstruction accuracy will slightly increase with a higher value of *K* (e.g. reconstruction accuracy is 0.69 for *K*=30). However, the instability score tends to decrease significantly for higher values of *K* (e.g. instability score for *K*=30 is 0.14 vs. 0.02 at *K*=11, which is roughly 7x higher). These findings indicate that staNMF outperforms PCA in automatically generating biologically-relevant patterns from spatial gene expression profiles.

### Gene expression-defined ontology from stable PPs

To draw connections between the PPs of gene expression and the mouse brain atlas, we investigated their overlap using spatial correlation analysis. We first calculated the Pearson correlation coefficient between all 868 expert-annotated brain regions from the Allen Common Coordinate Framework (CCF v3.0)^16^ to each of the staNMF PPs. We visualize 66 of the 868 regions in **Fig. 2** to facilitate the comparison. These 66 regions provide a complete medium-level representation of the mouse brain CCF. They are selected by including all “child” regions for the 12 coarse CCF regions (isocortex, olfactory areas, hippocampal formation, cortical subplate, striatum, pallidum, thalamus, hypothalamus, midbrain, pons, medulla, and cerebellar cortex/nuclei). In this paper, we define “coarse-level” regions as these 12 CCF regions, “medium-level” regions as their 66 children, and “fine-level” regions as all regions that are finer than medium-level.

**Figure 2:**
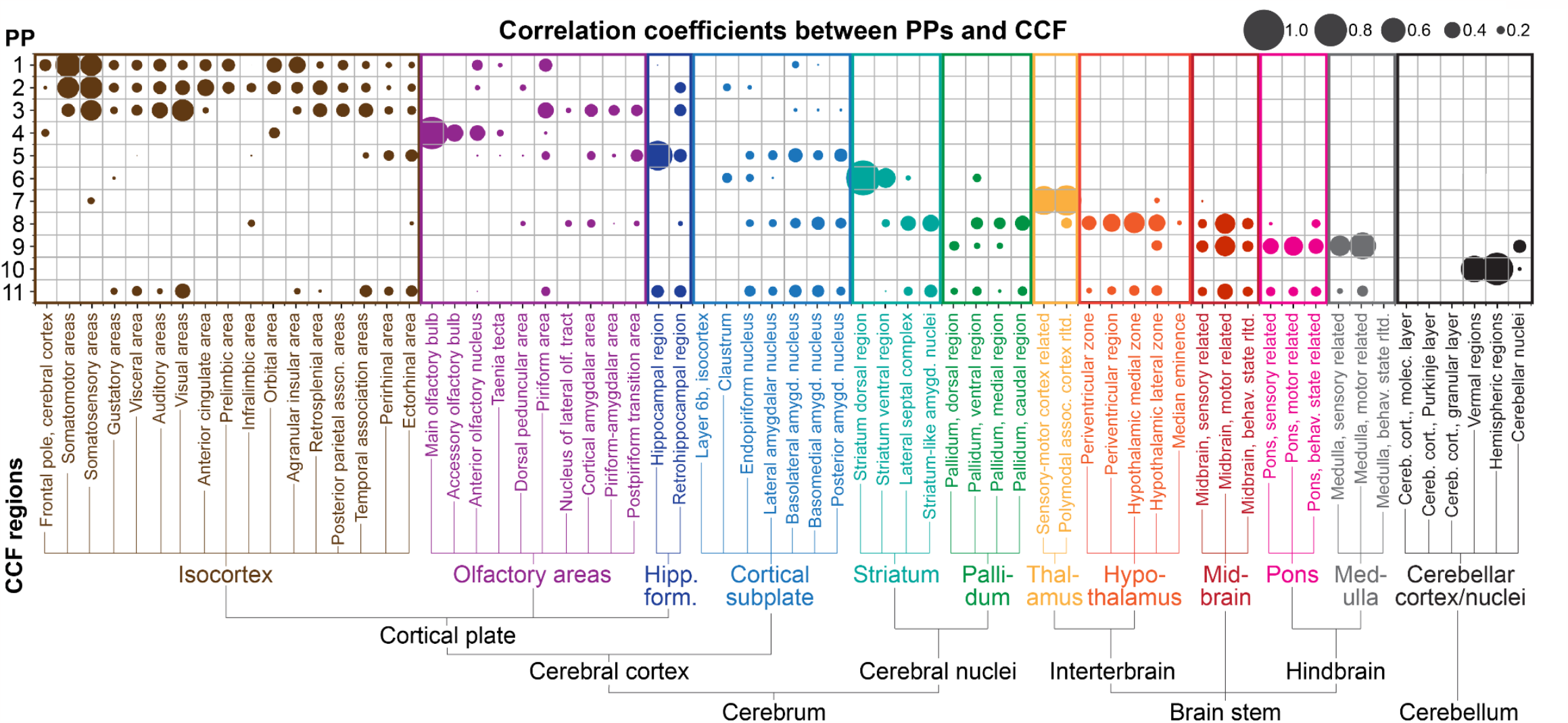
Region-dependent correlation between staNMF PPs and the CCF. Bubble plot of the Pearson correlation between PPs (y-axis) and expert-annotated regions from the CCF in the adult mouse brain (x-axis). Each bubble represents the value of the correlation coefficient between each PP and a CCF region. The CCF regions with labels are the complete set of 66 children of the 12 coarse CCF regions and are organized left-to-right based on the CCF ontology map. The PPs are organized top-to-bottom based on their Pearson correlation to the CCF coarse regions.

We found that the gene expression-defined PPs from staNMF have similarities to the CCF ontology, but also major differences (**Fig. 2**). Three PPs (PPs 1-3) are well, yet in many cases, differentially correlated with select parts of the isocortex: They all have correlation with the somatosensory areas of the isocortex, in addition to differential correlation with other cortical areas (e.g., somatomotor, visual, and orbital areas of the isocortex). Interestingly, PPs 1-3 also have varying representations outside of the isocortex, including in the olfactory areas, hippocampal formation, and cortical subplate, which are each viewed as part of the cerebral cortex^16^. PP4 is mostly represented within the olfactory areas, with an especially high correlation to the main olfactory bulb and orbitofrontal areas of the isocortex. PP5 has a strong correlation to hippocampal formation but has some correlations to sub-regions within the isocortex, olfactory areas, and cortical subplate. Thus, we see that PPs 1-5 correlate with the cerebral cortex, one of the three highest-level CCF regions (in addition to the brainstem and cerebellum), but do not fit neatly within the coarse- or medium-level CCF regions.

Moving next to PP6, we found a considerably high correlation between that and the striatum with minor expression in the cortical subplate. PP7 exhibits a high correlation only to the thalamus, showing good agreement with CCF’s thalamus in the overall ontology. Unlike PP7, PP8 shows correlations spread across multiple regions, especially the hypothalamus, midbrain, striatum, pallidum, and cortical subplate (in descending order of correlation), which suggests that these CCF regions share gene expression patterns. Similarly, PP9 is highly correlated with multiple regions in the brain stem areas including the medulla, midbrain, and pons, as well as a minor expression in cerebellar nuclei. PP10 is highly correlated with the cerebellum, with major expression in cerebellar vermal and hemispheric regions but not in the cerebellar nuclei. A comparison between PP9 and PP10 suggests that there are significant gene expression differences between the cerebellar nuclei and the vernal/hemispheric regions of the cerebellum. Genes that are expressed in cerebellar nuclei tend to also be expressed in the brain stem areas while genes that are expressed in cerebellar vernal/hemispheric regions tend to be exclusively present in the cerebellum. PP11 is correlated to most CCF regions and visual inspection (**Fig. 1C**) suggests that it corresponds to the noisy gene expression profiles throughout the brain.

Besides examining one-to-one relation between CCF ontology and PPs, we asked which combination of CCF regions is best aligned with each PP. To answer this question, we ran a search of all possible neighboring combinations of 2 or 3 CCF regions. From a total of 868 CCF regions, we found 22,711 binary combinations and 1,834,540 ternary combinations that are spatially contiguous. We did not consider higher-order combinations due to the exponentially growing search space. We then identified the maximum PCC between each PP and the superset of all single CCF regions, all combinations of 2 CCF regions, and all combinations of 3 CCF regions. We found that the PPs tend to align with combinations of the coarse, medium, and/or fine CCF regions, but these combinations may exhibit different ontology than CCF (**Fig. 3A**):

**Figure 3:**
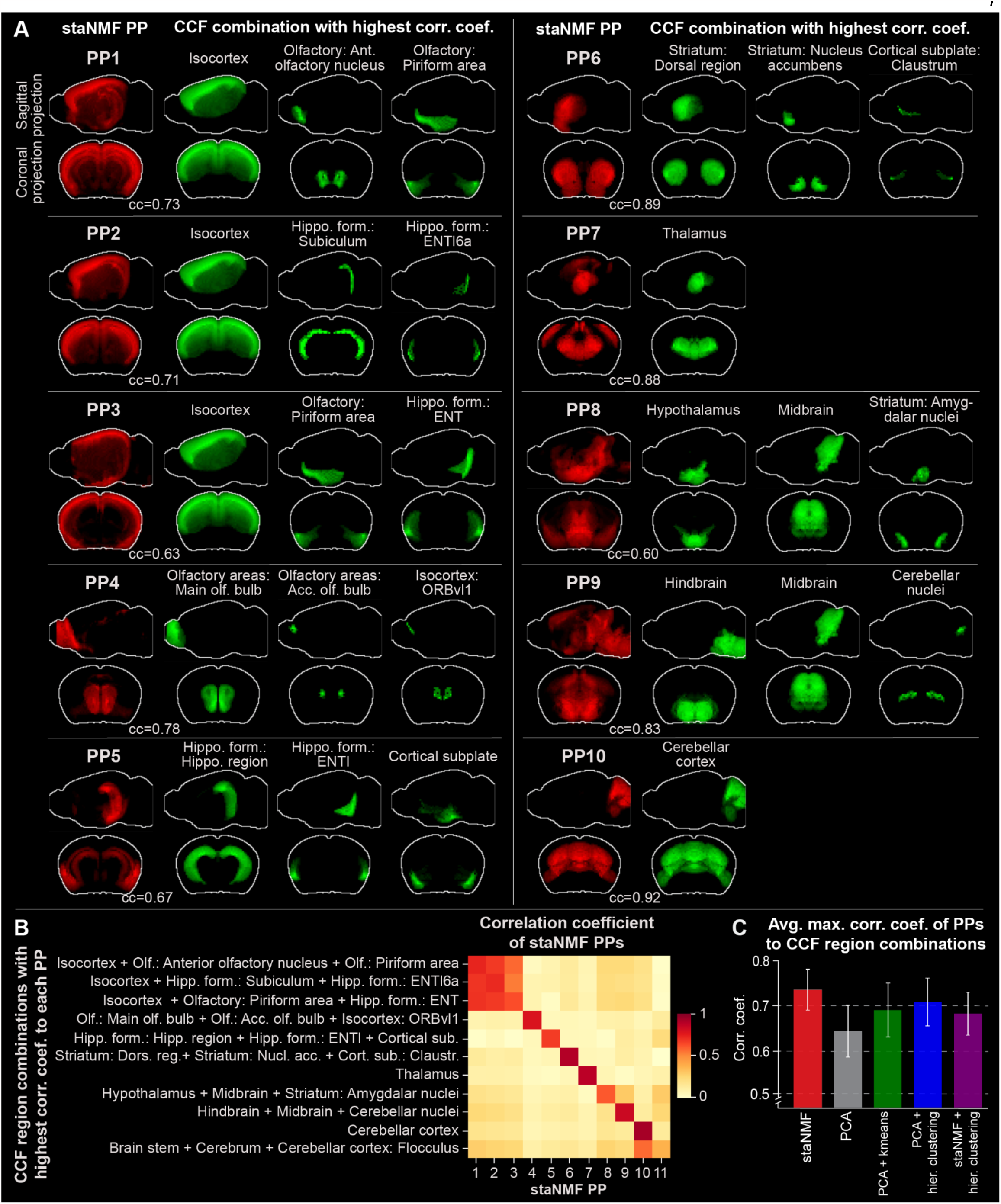
Alignment between staNMF PPs and combinations of CCF regions. **A**. PPs (in red) and the most similar combination of expert-annotated regions (in green) from the adult mouse brain CCF projected on the sagittal and coronal planes. The green regions are selected from a single or a combination of 2, or 3 neighboring regions from all 868 CCF regions with the highest PCC to each PP. The top 10 PPs are shown in descending order of the correlation coefficient. **B**. Heatmap of the correlation coefficient between staNMF PPs and each PP’s combination of CCF regions with the highest correlation coefficient. **C**. Comparison of the average maximum correlation coefficient of CCF region combinations to each PP for five matrix decomposition and segmentation methods: staNMF, PCA, PCA followed by k-means, PCA followed by hierarchical clustering, and staNMF followed by hierarchical clustering.

1. PPs 1-3 have their highest correlation to combinations of three CCF regions (correlation coefficients of 0.73, 0.71, and 0.63, respectively), which includes the isocortex. In addition to isocortex, PP1 adds the anterior olfactory nucleus and the olfactory piriform area, PP2 adds two finer-level retrohippocampal regions including the subiculum and the fine-level layer 6a of the lateral entorhinal area (ENTl6a), and PP3 adds the olfactory piriform area and the entorhinal area (ENT) of the retrohippocampal region.
2. PP4 has its highest correlation (correlation coefficient of 0.78) with the combination of olfactory bulb and accessory olfactory bulb with a fine-level cortical region (Orbital area, ventrolateral part, layer 1, referred to as ORBvl1).
3. PP5 is maximally correlated with a combination of two medium-level regions from hippocampal formation (hippocampal region and ENTl), and the high-level cortical subplate region.
4. PP6 has its highest correlation (correlation coefficient of 0.89) to a combination of three CCF regions: 1) striatum: dorsal region; 2) striatum: nucleus accumbens; 3) striatum: olfactory tubercle. PP6 does not include the striatum: amygdalar nuclei. Instead, the combination of hypothalamus, amygdalar nuclei, and midbrain is maximally correlated to PP8. Single-cell gene expression research has suggested that the amygdalar nucleus, midbrain, and hypothalamus contain cell types that are in fact highly related^54^.
5. PP7 and PP10 are the only PPs that are each maximally correlated with only one single CCF region: PP7 has a correlation coefficient of 0.88 to the thalamus, while PP10 has a correlation coefficient of 0.92 to the cerebellar cortex.
6. PP9 is maximally correlated with the combination of hindbrain, midbrain, and cerebellar nuclei (correlation coefficient of 0.84). It organizes the midbrain and hindbrain together, and suggests a relatively high similarity of gene expression between the midbrain, medulla, and pons, as observed with single-cell transcriptomics and clustering^54^.

These observations show that the PPs from gene expression partition the mouse brain differently than the CCF, suggesting a distinct ontology. We verify the uniqueness of the partitions by examining the correlation matrix between all PPs and their associated CCF combinations (**Fig. 3B**). Most PPs (except PPs 1-3, and 11) exclusively map to their associated CCF region combinations, suggesting low overlap between these PPs. The average maximum correlation coefficient between PPs and their respective CCF region combination is 0.74±0.04. By contrast, the average correlation coefficient between each PP and the CCF region combinations is 0.10±0.17, except for its highest correlation region. PP11 has the lowest maximum correlation coefficient (0.37, vs. 0.60 as the next lowest) with comparable correlation coefficients to other CCF regions, further suggesting its role in accounting for the noise in gene expression profiles. The findings from the spatial correlation analysis facilitates the construction of a gene expression-defined ontology based purely on spatial gene expression data (**Fig. S4**).

Subsequently, we conducted similar correlation analysis using the outcomes from common methodologies used in segmentation and clustering of spatial gene expression data. staNMF PPs have a higher average correlation coefficient to their respective CCF regions (0.73±0.05) compared to PCA PPs (0.63±0.06). Furthermore, the stronger diagonal pattern in the correlation matrix for staNMF (**Fig. 3B**) compared to PCA (**Fig. S3B**) suggests that staNMF PPs have a better alignment with the annotated brain regions. Additionally, we conducted the same spatial correlation analysis on PPs from typical clustering techniques. We clustered the ABA dataset using (1) PCA followed by k-means clustering (similar to the stLearn framework^55^), (2) PCA followed by agglomerative hierarchical clustering (similar to the AGEA framework^17^), and (3) staNMF followed by hierarchical clustering as a point of comparison (**Fig. 3C**). staNMF has the most similar PPs to their optimal CCF regions (correlation coefficient of 0.73±0.05), whereas PCA, PCA followed by k-means, and PCA followed by hierarchical clustering have lower correlation coefficients (0.63±0.06, 0.68±0.06, and 0.70±0.06, respectively). Additionally, staNMF PPs have a higher similarity to CCF region combinations compared to staNMF followed by hierarchical clustering (0.67±0.05). The comparison demonstrates that the PPs from staNMF alone are more similar to the combinations of known brain regions compared to PCA or standard clustering approaches.

### Substructures of mouse isocortex in PPs

The mouse isocortex is a layered structure^56^ with gene expression gradients along the anteroposterior and mediolateral axes^5^. This information, subject to the limit of data resolution, is also reflected in the PPs 1-3. We observe that for each of these PPs, the correlation coefficient to the isocortex dominates the correlation vs. the other regions that make up the highest correlated combination (**Fig. 4A**). For example, PP2 has a correlation coefficient of 0.70 to the isocortex, while it only has a 0.13 and 0.08 correlation coefficient to the other two regions that make up its highest correlated combination. Similarly, PPs 2 and 3 have correlation coefficients of 0.70 and 0.54 to the isocortex, respectively, while their correlation coefficients to other regions are considerably lower (**Fig. 4A**). Visualization of these three PPs suggests that they represent different spatial regions of the isocortex, in addition to minor components of the hippocampus and olfactory areas (**Fig. 4B**). PP1 represents the superficial layers in the frontal areas of the cortex, in addition to a partial representation of the anterior olfactory nucleus and the piriform area of the olfactory areas. PP2 represents the deeper layers of the isocortex in dorsolateral regions, and has a minor correlation to the subiculum and entorhinal area (lateral part, layer 6a) within the retrohippocampal region. PP3 represents the superficial layers of isocortex in dorsal regions as well as the piriform area of the olfactory areas and the entorhinal area of the retrohippocampal region. PP1 and PP3 have a gradual overlap in superficial layers, as indicated by the cyan color in **Fig. 4B**.

**Figure 4.**
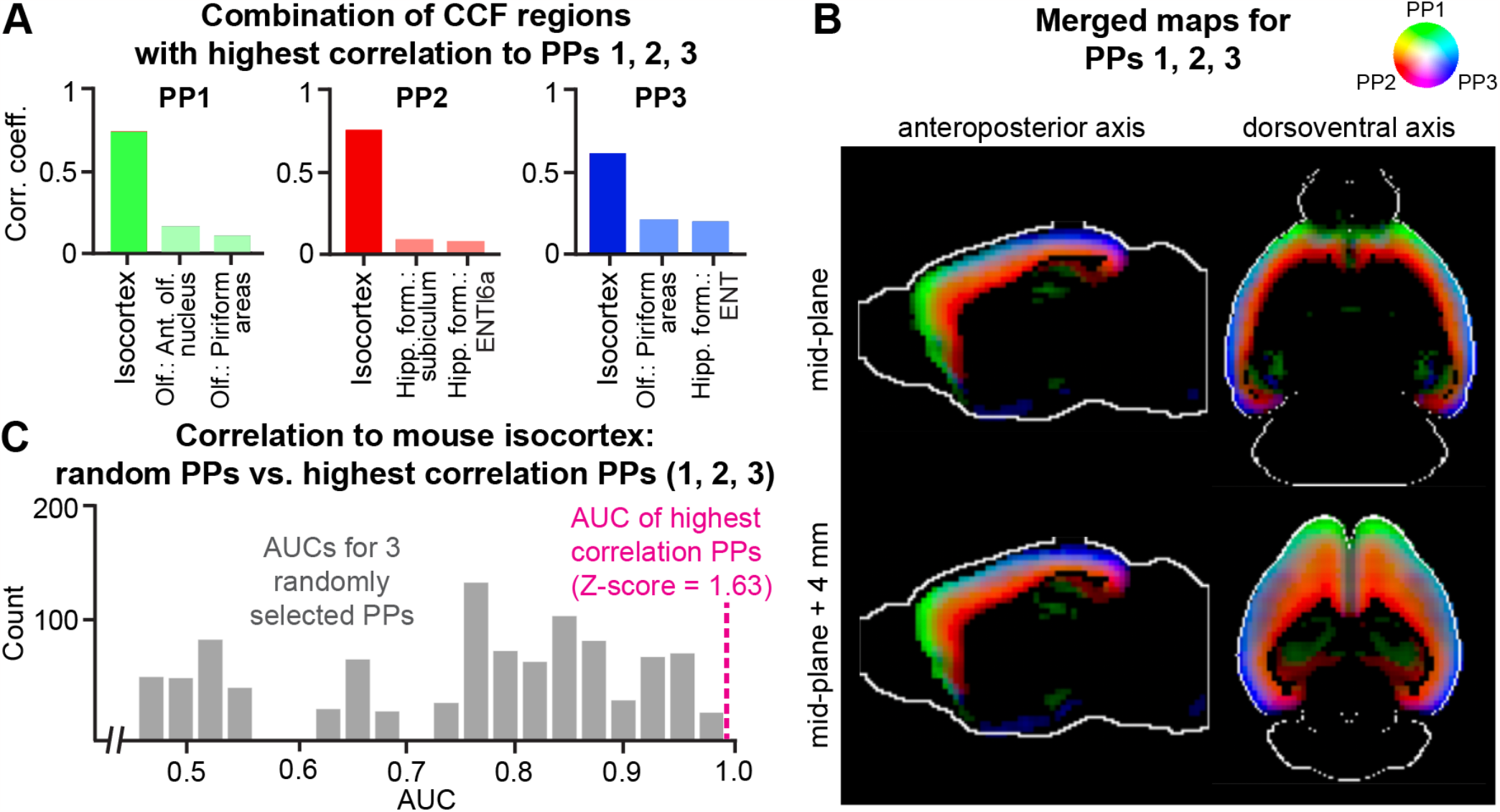
The defining PPs for the mouse isocortex. **A**. The three CCF regions with the highest correlations with PPs 1-3, with the isocortex being the most correlated region. **B**. Merged map of the PPs 1-3 in the isocortex. Each image is a 2D cross section viewed along the anteroposterior or dorsoventral axis (two columns). The rows represent the cross section used in visualization. **C**. Histograms of AUC values for isocortex for 1,000 runs of a logistic regression randomly fitting three PPs to isocortex CCF regions. The magenta vertical dashed line indicates that the AUC for PPs 1-3 are the best predictors of the isocortex compared to any other three random PPs.

We investigated how effectively the combination of PPs 1-3 can recreate the isocortex alone by training a logistic regression model to predict the isocortex CCF reference map from PPs 1-3. The area under the receiver operating characteristic (ROC) curve or AUC measure for the prediction is 0.99. The regression model is the most accurate model amongst 1,000 other models that uses a random selection of three PPs to predict isocortex (**Fig. 4C**). The median AUC for these 1,000 models is 0.78 (compared to 0.99 for the model that uses PPs 1-3, as shown in the magenta vertical dashed line), demonstrating that they represent the isocortex as a whole.

### From PPs to marker genes

Marker genes for an organ or tissue region are a set of genes with high expression within that region and relatively low expression in other regions. These genes are frequently used as starting points for understanding functions of cells and their local organization and to design genetic tools for experimental access to those cell types and regions for further knockout studies^57,58^. Given the relationship between PPs and brain regions established previously, one can robustly identify region-specific marker genes using the contributions of genes to the PPs. We visualize in **Fig. 5A** the gene-resolved coefficients (*a*_*kj*_) for each PP, where the genes are first ordered by the PP with the highest coefficient and then by their corresponding importance scores, *r*_*j*_*=a*_*kj*_*/*∑_*j*_ *a*_*kj*_ (*k=1,2*,…, *K*). The total number of genes selected by the PPs is not uniform across the board (**Fig. 5B**). Noticeably, PP5 (correlated with the hippocampal region) has by far the most unique genes, with over 1,500 genes. PP2 (correlated with the isocortex), PP9 (correlated with the hindbrain), and PP10 (correlated with cerebellar cortex) also have an especially large number of associated genes (represented by darker orange and red in the heatmap).

**Figure 5:**
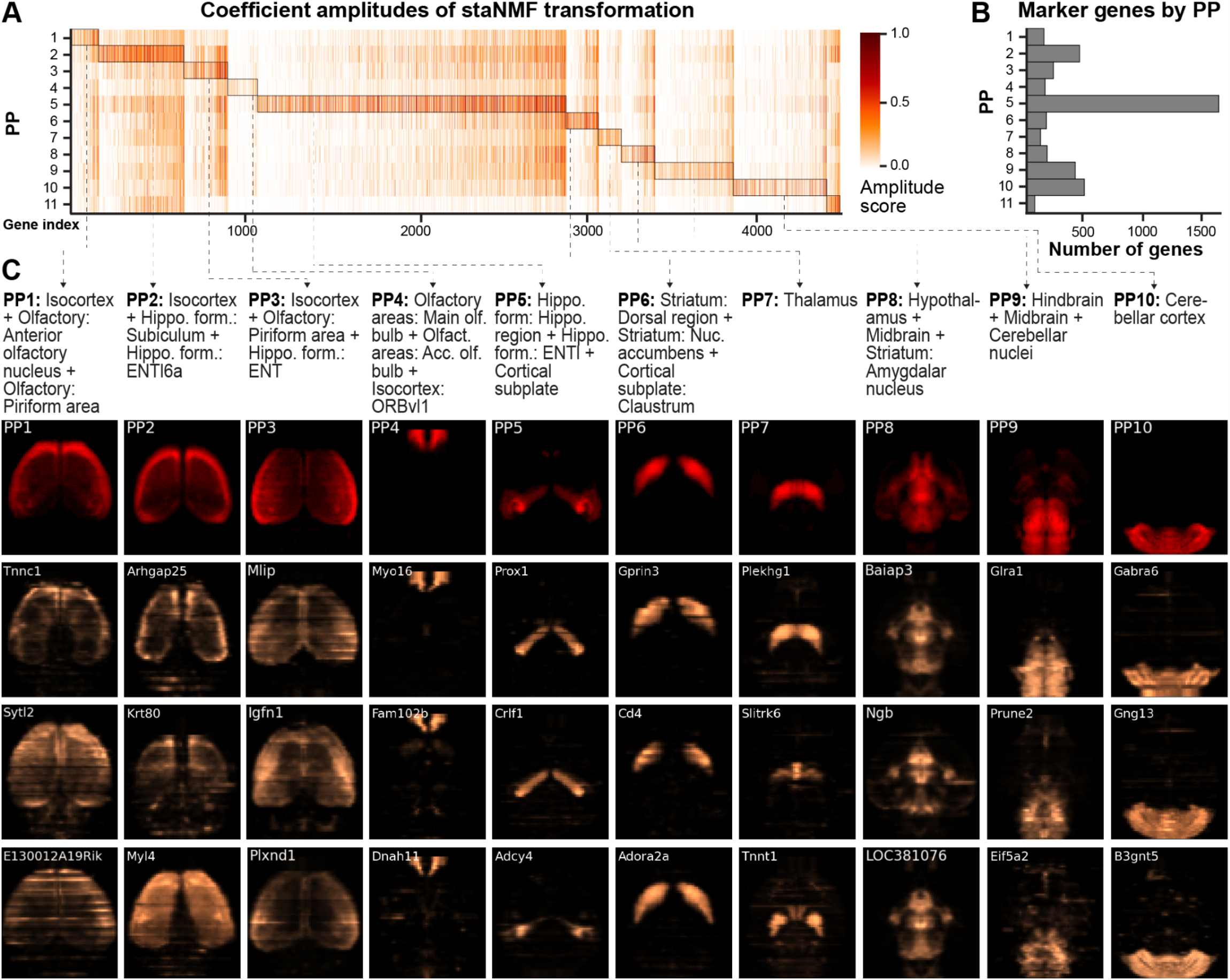
Marker gene identification from PPs. **A**. Heatmap of the gene-resolved coefficients of the 11 PPs, in descending order by the maximum correlation coefficient to a combination of CCF regions, as described in Fig. 3. Genes are assigned to the PP for which they have the highest coefficients. **B**. The number of significant genes for each PP, which is counted from the highlighted regions (by rectangular boxes) in A. **C**. The horizontal projection of the respective PP (in red) and the corresponding gene expression of the top three marker genes (in copper). The respective CCF region combinations are provided as text above. The name of the marker gene is displayed in the top left corner of each horizontal view of gene expression.

Drawing from these observations and the previous work on *Drosophila* embryos^18^, we used the following procedure to identify the marker genes for each PP: We first extracted the staNMF coefficients, {*a*_*kj*_} (*k=1,2*,…, *K*), for the *j*th gene. Each gene-resolved coefficient quantifies the contribution of the *k*th PP in explaining the expression of the *j*th gene. We then assigned each gene to a specific PP with the highest coefficient. Next, we calculate the importance score *r*_*j*_ for the *j*th gene and select the three genes with the highest ones for each PP as its associated marker genes. Consequently, the brain regions associated with the marker genes are determined, which shows convincing visual alignment (**Fig. 5C**). The regional designation of marker genes in the mouse brain has biological relevance. For example, *Prox1*, the top identified marker gene for PP5 (associated with hippocampal formation and cortical subplate), is known to be widely expressed across the brain during development, but primarily in the hippocampus and cerebellum in adulthood^59^. As another example, *Gabra6*, the top identified marker gene for PP10 (associated with the cerebellar cortex) is known to be preferentially expressed in the cerebellum as part of a program related to differentiation^60^.

### From PPs to spatial gene co-expression networks (sGCNs)

It is known that the spatial co-expression of genes yields meaningful biological relationships^61,62^. For example, an sGCN has successfully reconstructed the gap gene regulatory network in *Drosophila*^18^. However, few existing computational tools incorporate spatial information in identifying gene co-expression networks, and the ones that do only leverage existing expert-defined ontologies^63–66^. Data-driven ontologies from tools like staNMF will allow better identification and exploration of 3D spatial gene networks.

Building on a similar analysis for *Drosophila* embryos^18^, we used the following process to construct putative sGCNs for the PPs in the adult mouse brain: We first identified the top marker genes for each PP by selecting the genes with the top 0.025% importance scores correspondingly. Then, we computed the Pearson correlation coefficient between the staNMF coefficients of the two top marker genes of each PP. We drew an edge between two genes if their similarity score is among the top 5% of all similarity scores for that gene subset.

Our analysis resulted in the selection of 10 or 11 top marker genes for each PP, which are used to construct the putative sGCNs (**Fig. 6** for PPs 1-7, and **Fig. S5** for the remaining PPs). Interestingly, some of the regulatory relationships that are recently found via experimental research are present in the sGCNs. For example, in PP6, which is correlated to striatum, seven marker genes show especially strong edges (*Gprin3, CD4, Gpr6, Ric8b, Rgs9, Serpina9*, and *Gm261*) and seem to form a hub of connections. Interestingly, a 2019 experimental study in mice found that *Gprin3* controls striatal neuronal phenotypes including excitability and morphology, as well as behaviors dependent on the striatal indirect pathway and mediates G-protein-coupled receptor (GPCR) signaling^67^. *Gpr6* is a GPCR gene, and *Rgs9* and *Ric8b* are regulators of GPCR genes. In addition, *Gm261* and *Serpina9* are known to impact synapse development. In addition, *Prox1* and *PKP2* appear as interactions in PP5, which is related to hippocampal formation. Interestingly, a recently published experimental study has identified *Prox1* as a transcription factor associated with *PKP2* expression^68^. These relationships could be used as leads for experimental validation when studying specific genes in their tissue context.

**Figure 6:**
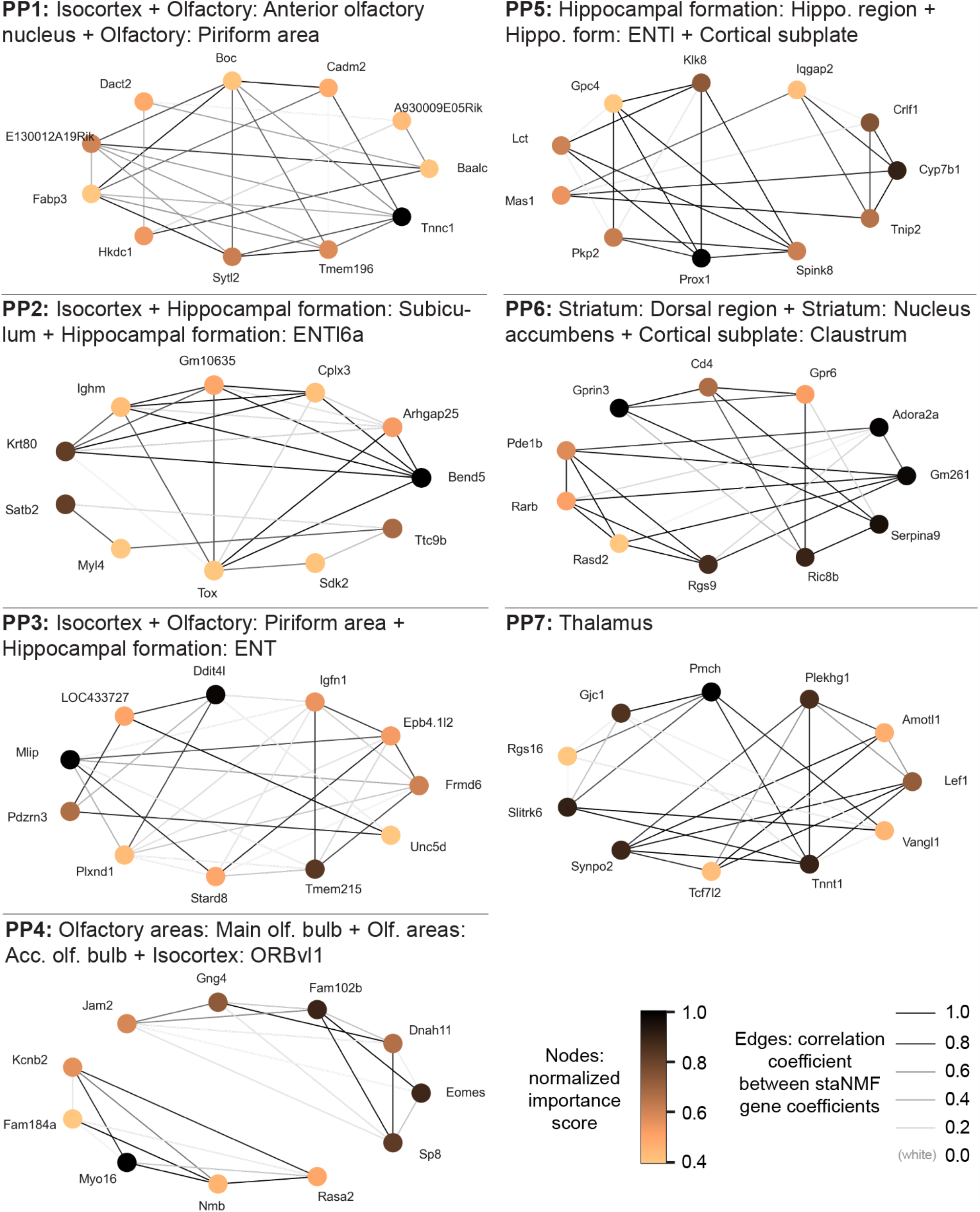
Putative spatial gene co-expression network (sGCN) construction. The sGCNs from PPs 1-7 and their associated brain regions from the CCF 3.0 are shown. The node color presents the selectivity of the gene to the PP associated with the brain region. An edge is drawn between genes if the similarity score is among the top 5% of all similarity scores for that gene subset. The edge thickness is proportional to the similarity scores between the staNMF representation of the two genes.

## Discussion

Unsupervised matrix factorization models are powerful machine learning tools for exploratory data analysis in spatial transcriptomics. Combining accurate unsupervised models with stable learning improves the interpretability of the resulting spatial patterns (i.e. PPs), as we have shown using staNMF in the present work. Our pipeline can automatically find consistent gene expression-defined spatial regions in 3D without supervision, eliminating the need for manual annotation. We point out here that anatomical atlases such as the ABA^16^ are constructed by dividing the brain volume into spatially contiguous regions without overlap. However, at the tissue or cellular level, this idealization is not always satisfied, and the strict division should be regarded as an approximation. Gene expression-defined PPs provide an automated way to explore whole-brain data with simplistic, spatially coherent regions that retain meaningful connections to the expert-annotated anatomical atlas.

Despite the limited spatial resolution, our analysis of the current dataset encompassing the entire adult mouse brain reveals promising marker genes with region specificity for future investigations in controlled experiments. As biological processes occur in 3D space and time, analysis of the co-expression network is inherently more reliable with data from 3D gene expression. The specific sets of genes and their spatial co-expression that contribute to PPs are also likely to contribute to the unique functions of brain regions they delineate. Those genes or their combinations identified through our spatial correlation analysis will be highly informative for designing genetic toolkits to experimentally access specific cell types and spatial domains within the organ of interest^69,70^. When combined with developmental data, these biological insights may help understand the longitudinal evolution of region-dependent gene expression to uncover signatures and functions hard to decipher from traditional transcriptomic methods without spatial information. Although the putative sGCNs identified here still need to be validated in controlled experiments, they may be linked to regulatory interactions, such as hub genes which are likely to mediate communication between networks, and to relationships between genes and gene modules. Our data-driven gene network representation might also be useful for studying disease processes such as selective vulnerability of certain regions to spread of pathogenic proteins in the brain^71^. Through integration with new scRNA-seq datasets, these networks can be used to study the cell-type specificity of spatial interactions between genes and find cell-type specific gene networks. The computational pipeline in the present work leverages the linear relationship between PPs to identify gene networks. Future work could incorporate nonlinear interactions using supervised methods such as iterative random forest^72^ to uncover complex gene interactions at the scale of the mouse brain.

Moreover, the availability of many different modalities for whole-organ imaging^6,7^ highlights the need for computational method developments along this direction. These methods would not only avoid human labor but are also more likely to be informative for investigating the functions of these regions. Besides gene expression data, the staNMF is also applicable to a broad range of biological data and may be used in multimodal data integration by combining learned representations. Potential future work will include integration with other modalities such as MRI and axonal projections to precisely characterize finer brain regions^56,73^. The computational efficiency of staNMF can be further improved to accommodate large datasets by exploiting the block structure of the data matrix or to use hierarchical updating schemes. The spatial neighborhood query in our computational pipeline may be upgraded into a discrete tree search to accommodate the existing brain ontology to explore higher-order combinations of brain regions. It is worthwhile to incorporate similar stability analysis in existing region-constrained matrix factorization models^39,40^ to assess the changes in the outcome. We are hopeful that the three principles for data science: predictability, computability, and stability (PCS)^4 7^ for veridical data science, as illustrated here, will be implemented in more case studies to improve the reproducibility of data-driven scientific discovery.

## Methods

### Data description and preprocessing

The primary dataset used in our study is the in situ hybridization (ISH) measurements from 4,345 genes at 200 μm isotropic resolution (a matrix size of 67×41×58 for each gene expression image) from the adult mouse brain at 56 days postnatal^48^. The data was collected at the Allen Institute for Brain Science and is publicly available under the Allen Brain Atlas (ABA) (https://mouse.brain-map.org/), as previously described^48^. An API enables the download of the data at (http://help.brain-map.org/display/mousebrain/API). The Allen Mouse Brain Common Coordinate Framework (CCF) was used as the 3D reference atlas^16^. We used CCFv3 which is publicly available at (http://help.brain-map.org/display/mousebrain/api), which consists of parcellations of the entire mouse brain in 3D and at 10 μm voxel resolution. CCF provides labeling for every voxel with a brain structure spanning 43 isocortical areas and their layers, 329 subcortical gray matter structures, 81 fiber tracts, and 8 ventricular structures. The methods for constructing the CCF dataset are previously described in detail^16^.

At preprocessing, we imputed missing voxels in the gene expression data using a k-nearest neighbors algorithm^74^ with 6 neighbors. To test the efficacy, we calculated the accuracy on a hold-out test set of 1,000 random voxels for each of the 4,345 genes from the ABA dataset (for a total of 4,435,000 data points). Following data imputation, we created a brain mask representing all the voxels of the mouse brain using the CCF which results in 55,954 voxels, vs. the total cube array of 159,326 voxels, reducing the number of voxels used for subsequent analysis by roughly two-thirds. Once the analysis was run, we unmasked the analysis outcomes and transformed the data back to the original shape (67×41×58). The data processing uses the codebase *osNMF* (https://github.com/abbasilab/osNMF), short for ontology discovery via staNMF.

### The staNMF framework

Non-negative matrix factorization (NMF)^32–34^ decomposes the data matrix into *K* dictionary elements and associated coefficients, resulting in parts-based representations of the original data. Stability-driven NMF (staNMF)^18^ is a model selection method that helps determine *K* through stability analysis. To apply staNMF to the 3D gene expression, we first transformed the imputed data into a matrix of voxels by genes (of size 55,954 by 4,345). The voxels were masked to leave out only those in the brain as previously described. The voxel-by-gene matrix is the input of the NMF algorithm which factorizes the gene data matrix into principal patterns (PPs). Formally, let **X** = [*x*_*1*_,*x*_*2*_,…, *x*_x_], be a *v* × *n* data matrix, where v is the number of unique voxels and *n* is the number of genes represented. Let **D** = [*d*_*1*_,*d*_*2*_,…, *d*_*k*_], be a *v* × *K* matrix, representing a dictionary with *K* elements or atoms (columns of **D**), and **A** = [*a*_*1*_,*a*,…, *a*_*n*_], be a *K* × *n* matrix, representing the coefficient matrix.

Under the current problem setting, NMF aims to minimize the loss function

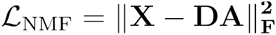

subjecting to non-negativity conditions **D** ≥ 0,**A** ≥ 0· The subscript F indicates the Frobenius norm. We used the *scikit-learn*^75^ implementation of NMF with default settings of the tolerance of the stopping condition (tol=0.0001) and the maximum number of iterations (max_iter=200). The staNMF is trained using coordinate descent (solver=‘cd’), which alternately optimizes the **D** and **A** matrices and is frequently used for NMF^76^.

The stability analysis for NMF selects the parameter *K* computationally using an instability score. In our work, we ran the NMF algorithm *N*=100 times at each integer value of *K* from 8 to 30. For each *K*, we compute an instability score that is the dissimilarity of learned dictionary pairs (**D** and **D**′) averaged over *N* runs. According to this definition, the optimal choice of *K* would result in highly stable dictionaries upon random initialization. The dissimilarity (dsim) is formulated using the cross-correlation (xcorr) matrix, **C**=xcorr(**D, D**′), between each dictionary pair and requires accounting for the scaling and permutation invariance of the learned dictionary elements^33,77^. Cross-correlation directly accounts for the scaling invariance between dictionaries in its normalization factor. To account for permutation invariance, we chose two distinct ways: The first way is to solve an assignment problem for the columns of **D** and **D**′ beforehand using the Hungarian matching (HM) method^49^, followed by calculation of the dissimilarity score,

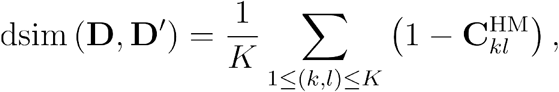

where (*k,l*) indicates assigned index pairs of **D**and **D**′ (*K* index pairs in total), the HM superscript indicates the cross-correlation matrix **C** calculated after applying the HM method. The second way is to account for the permutation invariance directly in the formulation of the dissimilarity metric using an Amari-type error function^50^,

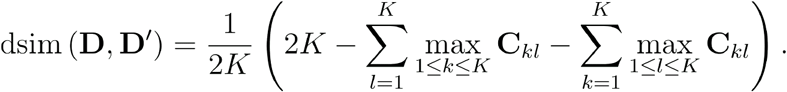

In either definition, the dissimilarity metrics are aggregated into the instability score γ using a simple average over *N*(*N*−*1*)*/2* distinct pairs^18^,

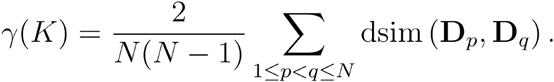

Stability analysis using either construction of the instability score yields the same result (*K* = *11*) for the most stable number of PPs (**Fig. 1B** and **Fig. S1**).

### Spatial neighborhood query

A brain atlas or parcellation *B*, with dimensions *a* × *b* × *c*, is a set of connected volumes, also called brain regions or parcels^15^, 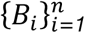 such that 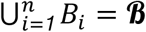. Numerically, each *B*_*i*_ is represented by a 3D segmentation mask, *B*_*i*_*∈ R*^*a*×*b*×*c*^, where the voxels within the mask (i.e. the support) have the value of *i* and those outside are 0. For the CCF 3.0 used in this work, *n=868*. The brain atlas is organized hierarchically based on biological knowledge of the brain regions, however, their precise spatial relationships are not explicitly given. We construct an adjacency list representation of the spatial relationship between brain regions for the subsequent analysis. This representation is commonly used in the spatial computing^78^ and image processing^79^ communities for its convenience. We call two brain regions, *B*_*i*_ and *B*_*j*_, are neighboring or spatially contiguous if they contain adjacent voxels. Because the support of each *B*_*i*_ has a different shape, we carried out the spatial queries of neighboring brain regions using image morphological (i.e. binary dilation) and logical operations to obtain the adjacency list. A pseudocode for generating all pairwise neighbors {(*B*_*i*_,*B*_*j*_)}of brain regions is given in **Algorithm 1**. The triplewise neighbors {(*B*_*i*_,*B*_*j*_,*B*_*l*_)}are generated similarly starting from existing pairwise neighbors, while the condition for spatial contiguity is that the third region (*B*_*l*_) after dilation is overlapping with at least one member (*B*_*i*_ or *B*_*j*_) of a neighboring pair.

#### Algorithm 1 Spatial neighborhood query (pairwise in 3D)

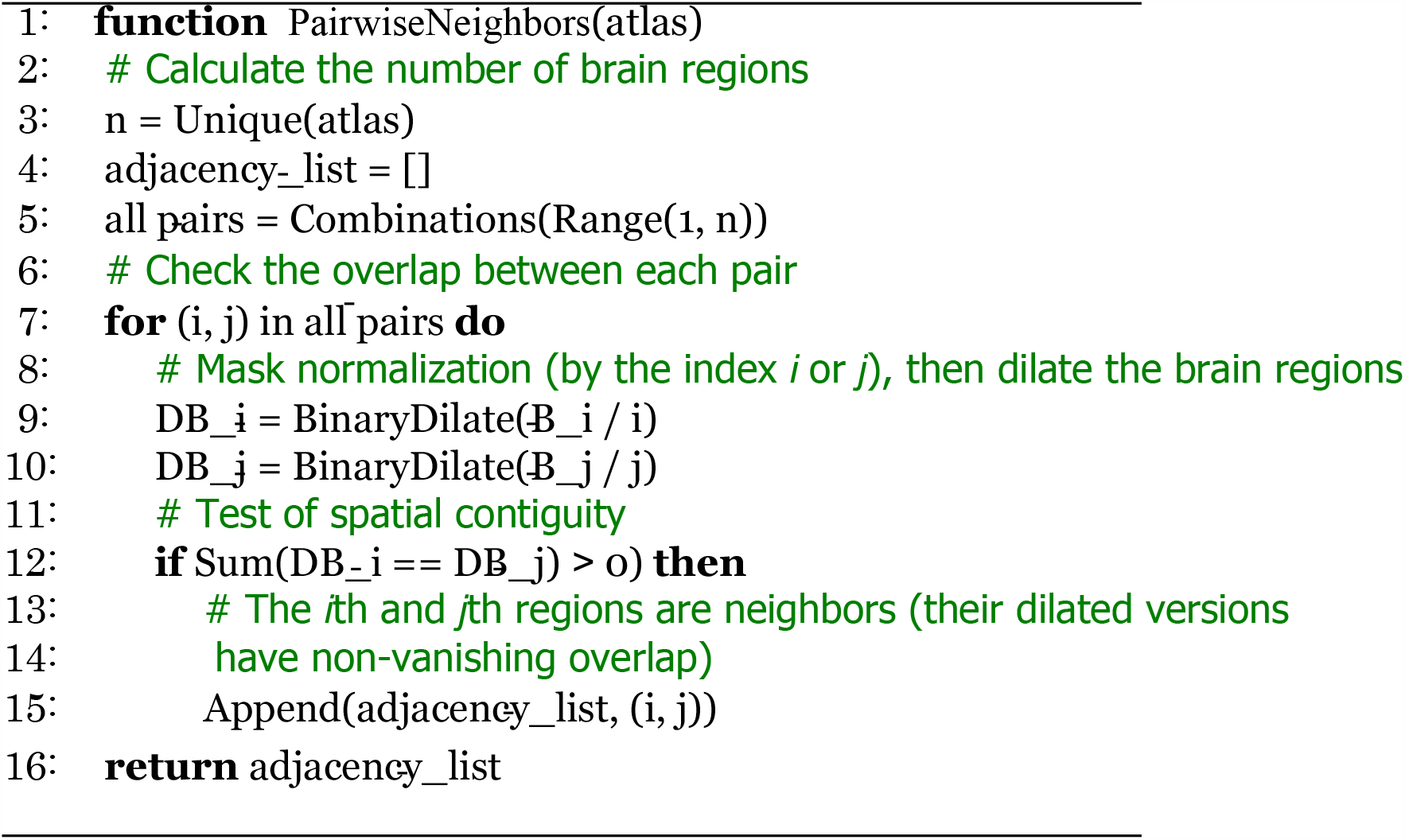

### Spatial correlation analysis

The overlap between the PPs and the brain regions are calculated by Pearson correlation (Corr). Let *X*_*k*_ be the volumes defined by a PP with index *k*, our spatial correlation analysis seeks the combination of spatially contiguous regions that maximize the Pearson correlation. For the *k*th PP, the expression for maximal correlation with two 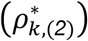 and three 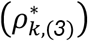 regions for are written as,

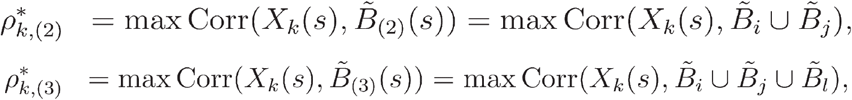

where 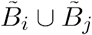 denotes the combined region of the pair (*B*_*i*_,*B*_*j*_) after mask normalization 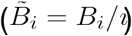. Similarly, 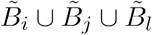 denotes the combined region of the triple (*B*_*i*_,*B*_*j*_,*B*_*l*_) after mask normalization. The terms 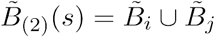 and 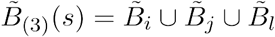 may be regarded as random variables indicating the random combinations of regions, where *s* denote the spatial coordinates. The maximization is conducted by exhaustive search over the respective adjacency list obtained from the spatial neighborhood search.

### Moran’s I

To quantify the spatial coherence of PPs, we used Moran’s I statistic^51^. It was originally used in geostatistics and has more recently been used in spatial gene expression literature^42,52,53^. Moran’s I ranges in value from -1 to 1. A value close to -1 indicates little spatial organization, similar to a chess board with black and white squares distributed across the board. A value close to 1 indicates a clear spatially distinct pattern, such as if all the black squares in a chess board were on one side and all white squares on the other. We calculated Moran’s I as follows^42^:

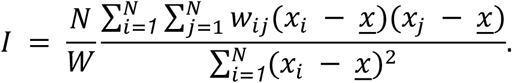

Here, *x*_*i*_ and *x*_*j*_ represent the PP coefficient at voxel locations *i* and *j*, respectively. *<u>x</u>* is the mean gene expression level of each PP. *N* is the total number of voxel locations, *w*_*ij*_ is the spatial adjacency relationship (based on the adjacency matrix, *w*) between voxels *i* and *j. W* is the sum of all entries in *w*, which represents the cumulative total adjacencies. We mask the dataset to only include the brain region. Then, for each voxel, we select up to 6 voxels for determining adjacency (up, down, left, right, forward, background, where available), following the “rook” definition of neighborhood. We assign *w*_*ij*_=1 if voxel *j* is adjacent to *i*, and *w*_*ij*_=0 otherwise. Given the large size of the adjacency matrix (159,326 × 159,326), we downsampled the PPs by removing every other row in each of the three dimensions to improve computational efficiency. Given certain voxels had multiple PPs with small but non-zero coefficients, we assigned each voxel in the brain map to the PP with the highest coefficient for that voxel. This ensures that unique voxels are not represented by multiple PPs.

### 3D visualizations of PPs

The 3D gene visualizations were performed using Napari viewer, a multi-dimensional image viewer for Python^80^. Key settings in Napari for PPs included: opacity=1, gamma=1, blending=‘additive’, depiction=‘volume’, and rendering=‘attenuated MIP’. MIP stands for maximum intensity projection, which enhances the 3D representation of objects. We moved the slide bar to 20% from the left side for ‘attenuated MIP.’

## Data availability

The data used in this study is publicly available under the Allen Brain Atlas (ABA) (https://mouse.brain-map.org/). The intermediate files are freely available at https://github.com/abbasilab/osNMF.

## Code availability

The software package is freely available at https://github.com/abbasilab/osNMF.

## Contributions

RA, YW, BY, BT, and HZ conceived the research design. RA and YW fitted the model and implemented the pattern discovery pipeline, marker genes analysis, and gene network reconstruction. RC and RA designed and implemented the ontology discovery pipeline and the spatial analysis. AJL implemented the visualization platforms. RPX, BY, BT, HZ, and AJL provided input on the analysis. RA and RC wrote the manuscript with inputs from RPX, BT, HZ, and BY.

## Acknowledgments

The authors would like to thank Lydia Ng and Zizhen Yao for their constructive feedback. RA, BT, BY, and HZ would like to acknowledge support from the Weill Neurohub through the Weill Neurohub’s Next Great Ideas Award. RA would like to acknowledge support from Sandler Program for Breakthrough Biomedical Research, which is partially funded by the Sandler Foundation.

## Supplementary Figures

**Figure S1:**
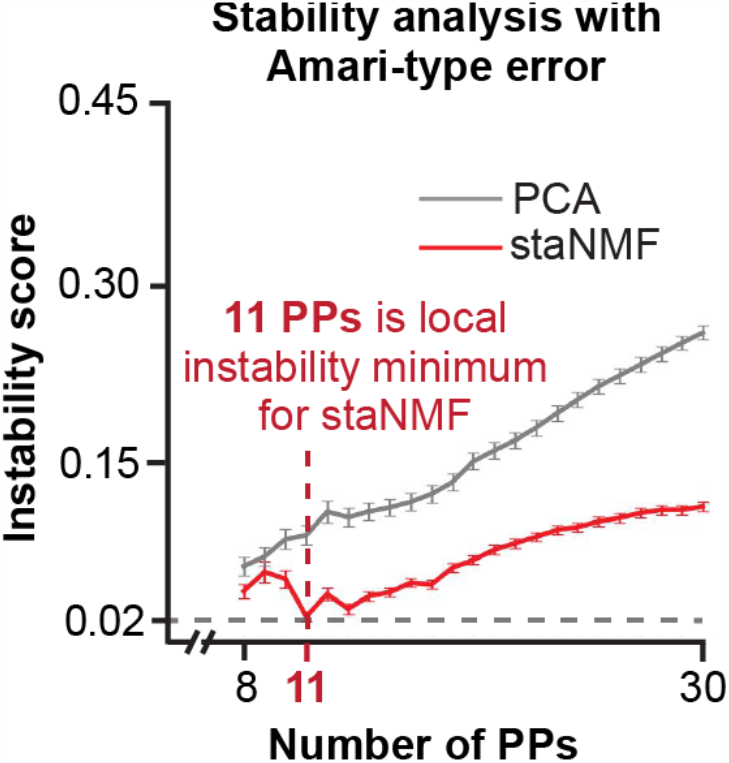
Stability analysis with Amari-type error function. Instability score of staNMF PPs and PCA PPs across 100 runs for each *K* value, from 8 to 30 for ABA dataset. The error bars are the standard deviation. This figure uses Amari-type error, while Fig. 1B uses the Hungarian matching method. Both approaches identify K= 11 for the minimum instability score (and thus most stability) for staNMF PPs.

**Figure S2:**
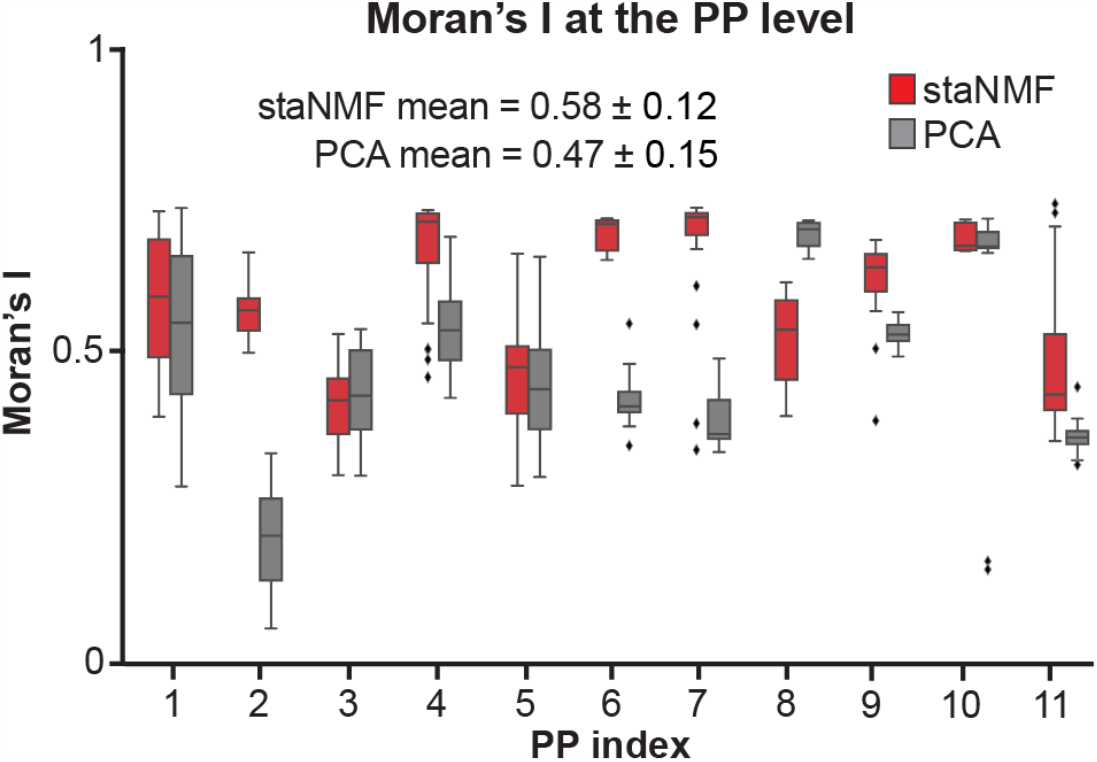
Moran’s I per PP from staNMF and PCA. The plot uses data from 20 bootstrapped simulations for each PP, for a total of 220 simulations for staNMF PPs and 220 simulations for PCA PPs. The mean Moran’s I was 0.58 ± 0.12 for staNMF and 0.47 ± 0.15 for PCA. The p-value between the two samples was <0.001. The PPs from staNMF show greater spatial coherence, or higher Moran’s I, than those from PCA for all but one case (PP8).

**Figure S3:**
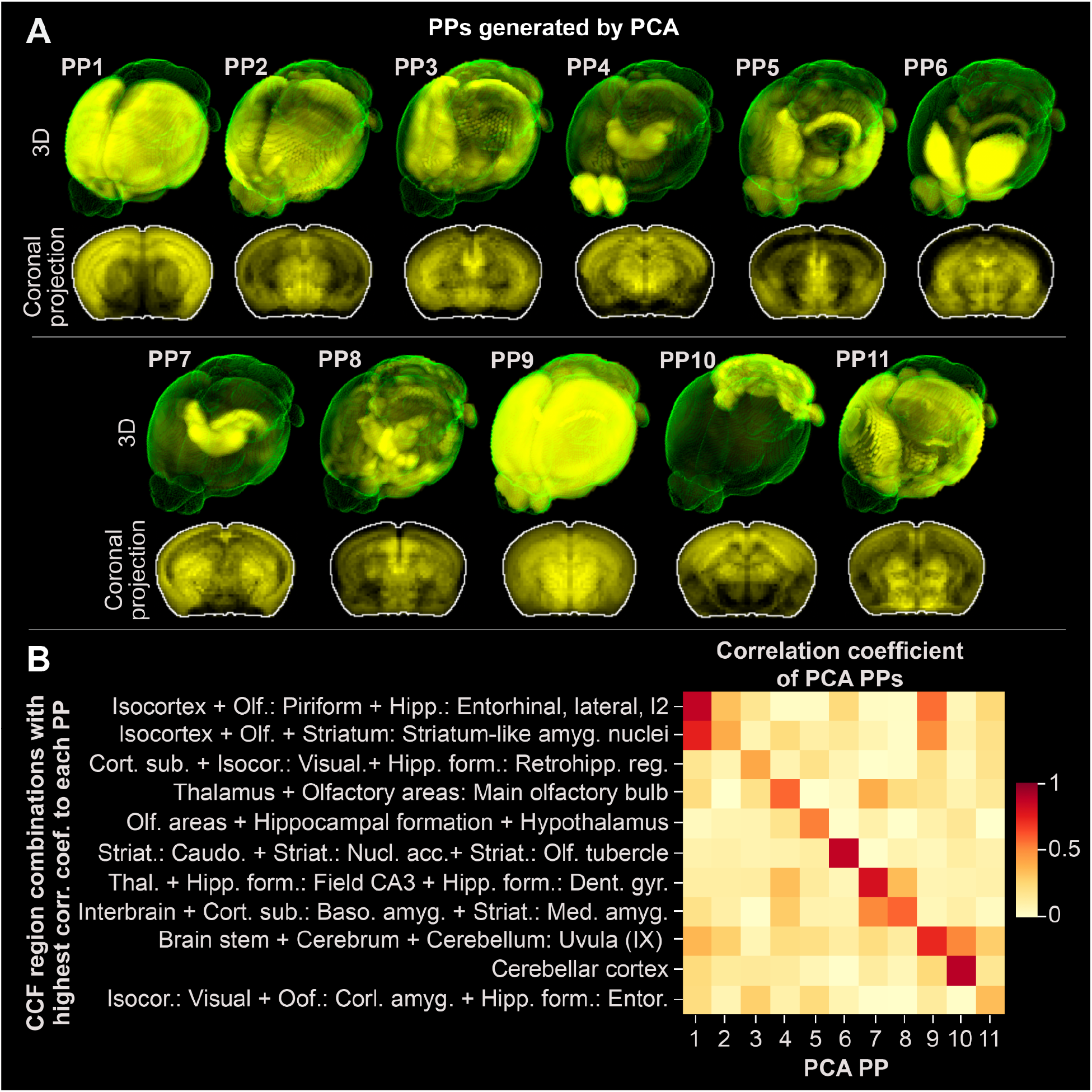
Similarity of PCA PPs to the expert-annotated brain regions. **A**. 11 PPs generated by PCA, ordered based on highest coarse region correlation to CCF ontology in 3D and projected on the coronal plane. **B**. Heat map of the correlation coefficient between PCA PPs and the most similar combination of CCF regions (with the highest correlation coefficient).

**Figure S4:**
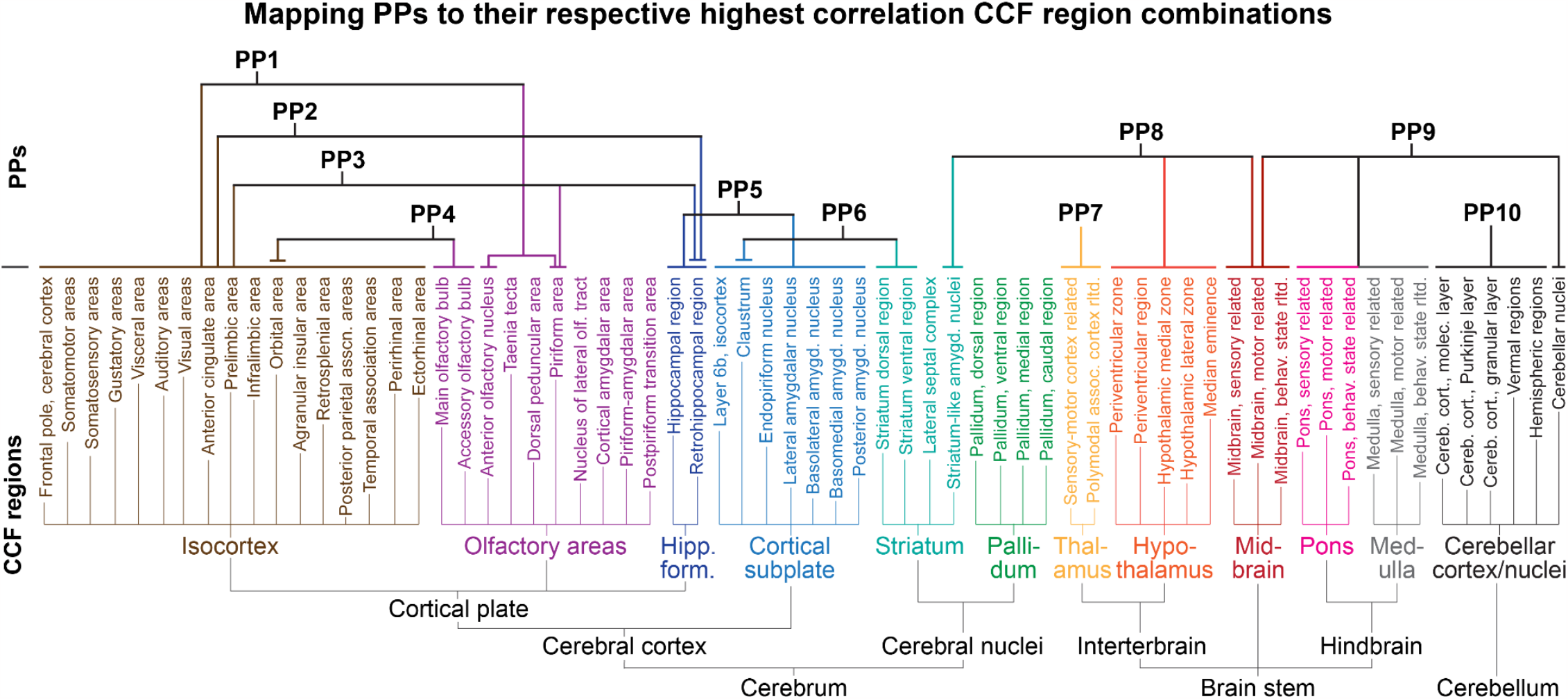
Summary of staNMF PPs mapping to the best-fit combination of Allen CCF regions. The 10 PPs from main text Fig. 3 mapped to their best-fit combinations of CCF regions.

**Figure S5:**
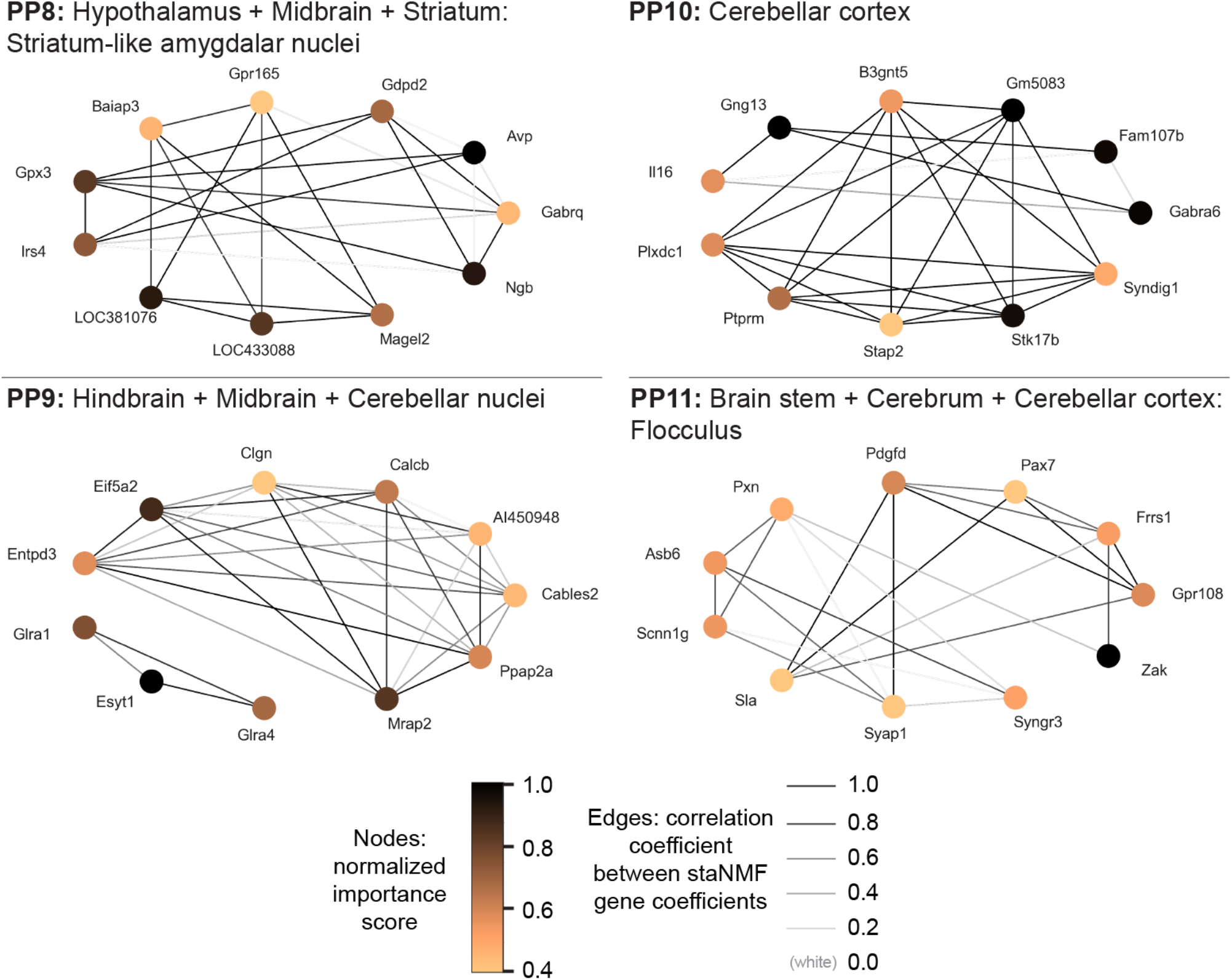
Putative spatial gene co-expression network construction, continued. Extension of Fig. 6 of the main text, the sGCNs from PPs 8-11 and their associated brain regions from the CCF 3.0 are shown. The node color presents the selectivity of the gene to the PP associated with the brain region. An edge is drawn between genes if the similarity score is among the top 5% of all similarity scores for that gene subset. The edge thickness is proportional to the similarity scores between the staNMF representation of the two genes.

